# Evolution of g-Type Lysozymes in Metazoa: Insights into Immunity and Digestive Adaptations

**DOI:** 10.1101/2024.10.03.616589

**Authors:** Krishanu Mukherjee, Leonid L. Moroz

## Abstract

Exploring the evolutionary dynamics of lysozymes is critical for advancing our knowledge of adaptations in immune and digestive systems. Here, we characterize the distribution of a unique class of lysozymes known as g-type, which hydrolyze key components of bacterial cell walls. Notably, many basal metazoan groups, including ctenophores, sponges, and choanoflagellates (the sister group of Metazoa), lack g-type lysozymes. We reveal a mosaic distribution of these genes, particularly within lophotrochozoans/spiralians, suggesting the lateral gene transfer (LGT) events from predatory myxobacteria played a role in their acquisition, enabling specialized dietary and defensive adaptations. We further identify two major groups of g-type lysozymes based on their widespread distribution in gastropods. Despite their sequence diversity, these lysozymes maintain conserved structural integrity that is crucial for enzymatic activity, underscoring independent evolutionary pathways where g-type lysozymes have developed functionalities typically associated with different lysozyme types in other species. Specifically, using *Aplysia californica* as a reference species, we identified three distinct g-type lysozyme genes: two are expressed in organs linked to both feeding and defense, and the third exhibits broader distribution, likely associated with immune functions. These findings advance our understanding of the evolutionary dynamics shaping the recruitment and mosaic functional diversification of these enzymes across metazoans, offering new insights into ecological physiology and physiological evolution as emerging fields.

## Introduction

Lysozymes, first discovered by Alexander Fleming in 1922, are key antimicrobial enzymes within the innate immune system, widely known for their role in hydrolyzing the 1,4-beta-linkages between N-acetylmuramic acid (NAM) and N-acetyl-D-glucosamine (NAG), which are critical components of bacterial cell walls. This enzymatic activity is ubiquitous across various biological systems and tissues, including mammalian blood, tears, sweat, and milk, as well as in the egg whites of birds (Fleming, 1922; Rupley, 1967). Beyond their enzymatic role, lysozymes also exhibit potent antibacterial properties, as even their proteolytic fragments demonstrate antimicrobial effects (Ibrahim, Matsuzaki, & Aoki, 2001).

Lysozymes are classified into several distinct types, each with evolutionary variations. These include the c-type (chicken-type), g-type (goose-type), i-type (invertebrate-type), and the less common ch-type (*Chalaropsis*-type). Additionally, lysozymes are found in bacteriophages (phage-type), bacteria (bacterial-type), and plants (plant-type), each contributing to breaking down bacterial cell walls and immune defense in different organisms. Despite differences in their tissue and organismal origins, and minimal sequence similarity, lysozymes exhibit conserved protein crystal structures, indicating convergent evolution across eukaryotic kingdoms from animals to plants and even prokaryotes (Beintema & Terwisscha van Scheltinga, 1996; Callewaert & Michiels, 2010; Fischetti, 2005; Grutter, Weaver, & Matthews, 1983; Holtje, 1996; Monzingo, Marcotte, Hart, & Robertus, 1996; Weaver et al., 1984).

In vertebrates, lysozymes are key components of the innate immune system, serving as a frontline defense against invading pathogens. These enzymes are typically associated with immune cells such as monocytes, macrophages, and neutrophils (Ragland & Criss, 2017; Rosowski, 2020). Neutrophils, in particular, are produced in the bone marrow and circulate in the bloodstream, mobilizing quickly during infections (Burgener & Schroder, 2020). These neutrophils are major phagocytic cells across vertebrates, including fish, amphibians, reptiles, birds, and mammals. During immune responses, neutrophils perform key functions such as phagocytosis, apoptosis, and degranulation, and can form neutrophil extracellular traps (NETs) to capture and neutralize pathogens (Brinkmann et al., 2004; Fingerhut, Dolz, & de Buhr, 2020).

While vertebrate lysozymes are well-studied, little is known about highly diverse invertebrate groups, many of which rely entirely on innate immunity due to their lack of adaptive immune systems. In invertebrates, lysozymes serve both defense and digestive functions, particularly in filter-feeding species such as bivalves. These enzymes help protect against pathogens and can digest bacteria filtered through their gills. This dual role of lysozymes highlights a significant evolutionary adaptation in invertebrates, where digestive and immune functions have co-evolved together. Specifically, arrays of i-type lysozymes play critical roles in these processes, contrasting with the c-type lysozymes’ recruitment in vertebrate ruminants, where they facilitate similar digestive functions (Itoh, Okada, Takahashi, & Osada, 2010; Itoh & Takahashi, 2009; Matsumoto, Nakamura, & Takahashi, 2006).

The understanding of g-type lysozymes in invertebrates remains limited. These lysozymes were originally discovered in the egg whites of the *Anser anser* (Embden goose) and have since been identified in other chordates as well as various molluscs, such as scallops and mussels. This widespread presence across different phyla suggests that g-type lysozymes perform essential functions in both ecological and evolutionary contexts. Their functions range from aiding in digestion to providing defense against pathogens, highlighting their versatility and significance in the immune responses of these organisms (He et al., 2012; Hikima, Minagawa, Hirono, & Aoki, 2001; Nilsen, Myrnes, Edvardsen, & Chourrout, 2003; Wang, Zhang, Zhao, You, & Wu, 2012; S. H. Zhang et al., 2012; Zhao et al., 2007; Zou, Song, Xu, & Yang, 2005).

Lateral Gene Transfer (LGT) or Horizontal Gene Transfer (HGT) plays a significant role in biological innovations, shaping the evolutionary trajectories by cross-kingdom signaling and transfer novel genetic material across species. Recent studies have highlighted the importance of HGT in enabling organisms to acquire adaptive traits such as enhanced metabolism, versatile immunity, and reproductive functions (Li et al., 2022; Liu et al., 2023). The identification of g-type lysozymes in metazoans, which exhibit strong sequence similarities to bacterial lysozymes, particularly those of predatory myxobacteria, suggests that HGT may have contributed in their acquisition. This process may explain the mosaic distribution of g-type lysozymes across taxa, as observed in our study.

Here, we conducted an extensive screening of g-type lysozymes across metazoans, uncovering a complex and intermittent distribution pattern, particularly within the lophotrochozoan/spiralian superclade. Our findings suggest that this distribution may be driven by lateral gene transfer (LGT) events from bacteria. We also identified two major g-type lysozyme groups in gastropods, using *Aplysia californica* as the reference species. Notably, our analysis suggests that *Aplysia californica* and kin lacks both c-type and i-type lysozymes, indicating its reliance on three unique g-type lysozymes. Of these, two are highly expressed in the hepatopancreas, a key digestive organ, aligning with findings from other mollusks. The third g-type lysozyme shows variable expression across organs associated with both feeding and immune defense, highlighting the gene’s functional diversity.

These targeted expression patterns point to evolutionary adaptation through convergent evolution and lateral gene transfer to meet specialized dietary and defense challenges and needs. Additionally, we observed high sequence similarity between metazoan g-type lysozymes and those in myxobacteria, suggesting horizontal gene transfer from bacteria to animals. These findings provide insights into the evolutionary dynamics of lysozymes, shedding light on their mosaic functional diversification across metazoans.

## RESULTS

### Mosaic Distribution and Evolutionary Insights of g-Type Lysozyme Genes Across Metazoans

Two copies of g-type lysozyme (Lyg1 and Lyg2) have been identified in all extant mammals in a study that analyzed 250 species, with the exception of cetaceans and sirenians, which have lost both copies. Additionally, a potential loss of these genes was observed in the tarsier (X. Zhang et al., 2021). Similarly, genome sequences of two bony fish species (gar and tilapia) revealed no g-type lysozyme genes, while other teleost fish species possessed single or multiple g-type lysozyme genes (Irwin, 2014). Interestingly, g-type lysozyme genes have been found in various bird species, though not universally, as some species, like ducks, may carry pseudogenes (Irwin, 2014).

Considering the wealth of data available on vertebrates, we chose to shift our focus beyond vertebrates to explore the presence of g-type lysozyme genes in other lineages.

To further explore the evolutionary presence of g-type lysozyme genes, we performed an extensive screening for the presence of genes encoded g-type lysozymes across the sequenced genomes from representatives of all five basal metazoan lineages: Ctenophora, Porifera, Placozoa, Cnidaria, and Bilateria, viewing ctenophores as the sister lineage to the rest of Metazoa (Moroz et al., 2014; Ryan et al., 2013; Schultz et al., 2023; Whelan, Kocot, Moroz, & Halanych, 2015; Whelan et al., 2017). We expanded our investigation to include choanoflagellates and other holozoans, as well as prokaryotes, encompassing single-cell eukaryotes, protists, and fungi. In this survey, by screening five genomes (*Pleurobrachia bachei, Hormiphora californensis, Bolinopsis microptera, Mnemiopsis leidyi*, and *Beroe ovata*, see (Moroz et al., 2014; Ryan et al., 2013; Schultz et al., 2023; Vargas et al., 2024) and dozens of transcriptomes (Moroz et al., 2014; Whelan et al., 2017), we were not able to identify recognizable g-type lysozymes in ctenophores. In the homoscreromorph poriferan *Corticium candelabrum* we found three g-type related lysozymes, which we used as an outgroup, and no similar genes in the other three major groups of sponges (demosponges, glass, and calcareous sponges). The genomes of two haplotypes of *Trichoplax* (Placozoa) contain just one g-type lysozyme gene (Figure 1). Importantly, the genomes of both *Trichoplax* and *Corticium candelabrum* lacked other lysozyme types, such as c-type or i-type. We also discovered a single g-type lysozyme gene in the soft coral species *Xenia sp*. and *Dendronephthya gigantea* (Figure 1). These genes were not found in other cnidarian classes, such as hydrozoans and anthozoans. Notably, that a single g-type gene identified in brachiopods is nested within chordates. *Branchiosoma floridae* (lancelet, Cephalochordata) has six g-type lysozyme genes, one of the largest species-/genus-specific radiation events across metazoans.

**Figure 1:**
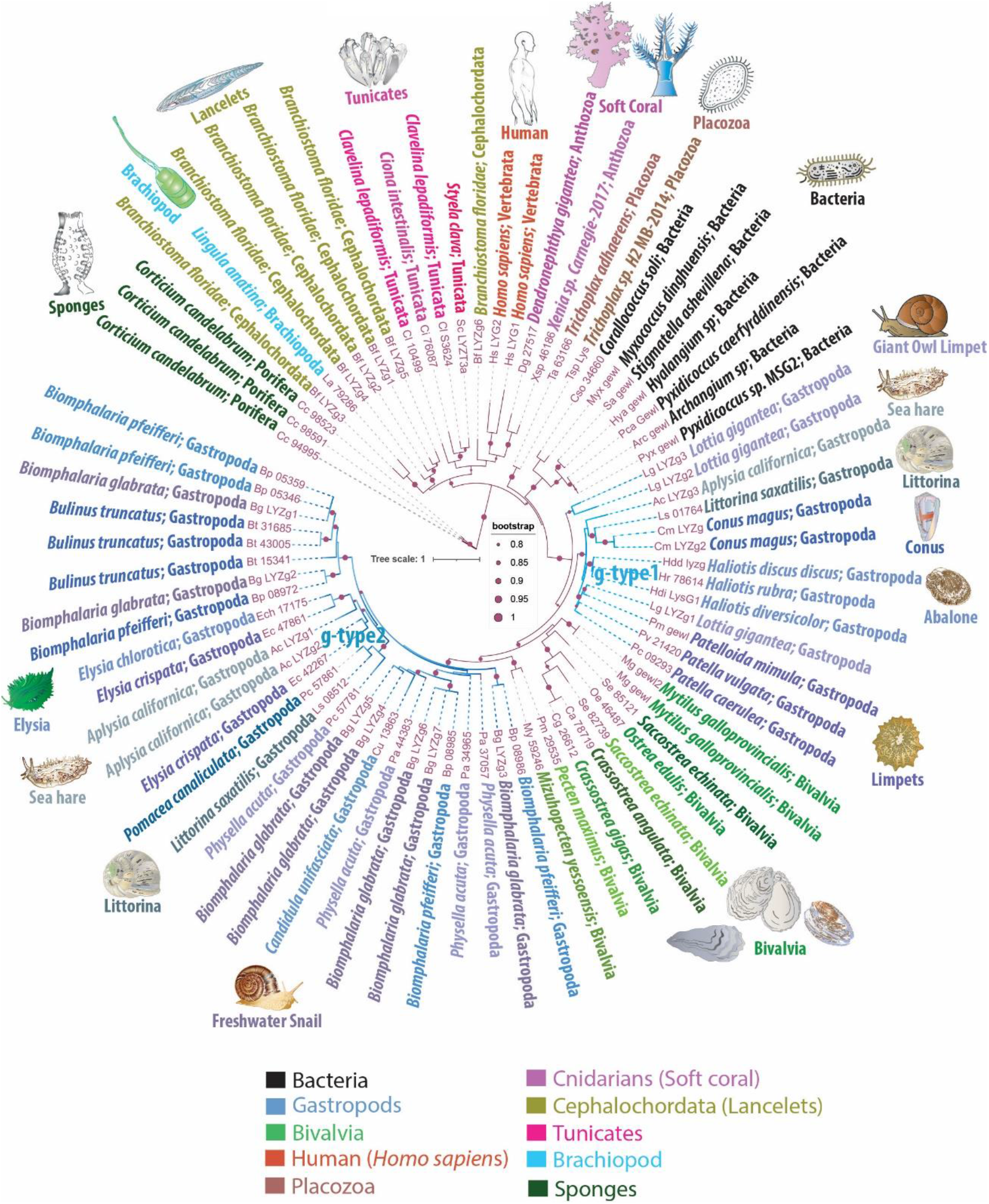
Phylogenetic Tree Illustrating the Mosaic Distribution of g-Type Lysozyme Genes Across Metazoans. This phylogenetic tree depicts the evolutionary mosaic distribution of g-type lysozyme genes across animal taxa, with gene names displayed alongside their respective genus and species name and the taxonomic group they belong to. Bootstrap values below 80% have been excluded for clarity. Lysozymes from predatory myxobacteria, highlighted in black, exhibit strong sequence similarities with metazoan g-type lysozyme genes, suggesting potential horizontal gene transfer events. The tree distinguishes two groups of lysozymes: g-type1, representing more ancestral genes found in basal species of gastropods, and g-type2, which appears to have arisen from more recent gene duplication events and is primarily found in the more recently evolved *Euthyneura* group, including species like *Elysia spp*., *Biomphalaria spp*., *Physella acuta*, and *Littorina saxatilis*.

This sporadic presence and distribution patterns strongly suggest that basal metazoan lineages acquired the g-type lysozymes via LGT events. We observed phyletic relationships between animal g-type lysozyme genes and several bacterial genes (Figure 1), illustrating the robust branching of prokaryotic genes within animal g-type lysozyme genes.

Notably, the sequenced genomes of five unicellular holozoans, sisters to Metazoa (*Monosiga brevicollis, Capsaspora owczarzaki, Pigoraptor chileana, Pigoraptor vietnamica*, and *Ministeria vibrans*), do not contain g-type lysozyme genes. However, these holozoan genomes contain molecularly distinct, unclassified lysozyme genes that do not belong to the traditionally recognized types, such as i-type or c-type lysozymes (KM and LLM, manuscript in preparation). Similarly, our screening of fungal genomes also did not yield any g-type lysozyme genes. Moreover, phylogenetic analyses revealed significant sequence homology between animal g-type lysozyme genes and those found in several bacterial species, as depicted in Figure 1. These bacterial homologs, belonging to myxobacteria, include representatives of the genera *Corallococcus, Myxococcus, Pyxidicoccus, Archangium, Stigmatella*, and *Hyalangium*. These myxobacteria are well-known for their *predatory behaviors*, utilizing complex mechanisms to hunt and consume other microorganisms, including bacteria, fungi, and algae (Contreras-Moreno, Perez, Munoz-Dorado, Moraleda-Munoz, & Marcos-Torres, 2024). This phylogeny implies horizontal gene transfer between these predatory bacteria and metazoans, underlining the adaptive significance of g-type lysozymes for innate immunity and defense, and early possible metazoan diversification.

The molluscs, and gastropods, in particular, revealed the extensive acquisition and lineage-specific radiation of g-type lysozymes. The graser, freshwater snail *Biomphalaria glabrata* has seven g-type genes and five lysozyme-encoded genes present in closely related species of *Biomphalaria pfeifferi*. Most other gastropod species have 2-4 copies of g-type lysozyme genes per genome (Figure 1). One or two copies of g-type lysozymes were found in the sequenced bivalves’ genomes. In contrast, cephalopods, *Octopus bimaculoides, Octopus vulgaris, Octopus sinensis*, and *Amphioctopus fangsiao*, each possess one or two i-type lysozyme genes, but none had g-type lysozymes. We did not identify any g-type lysozymes in the sequenced annelid, nemertines or phoronid genomes.

In summary, the distribution patterns of g-type lysozyme genes suggest two possible evolutionary scenarios: either the g-type lysozyme gene was initially present in the common ancestor of all metazoans and subsequently underwent widespread losses throughout ∼530 million years of subsequent evolution, or certain species sporadically acquired it within some metazoan lineages through domestication or horizontal gene transfer. The mosaic distribution and the rare occurrence of g-type lysozyme genes rather support the latter scenario. This indicates a selective evolutionary process characterized by intermittent gene retention, loss, and horizontal gene transfer within this particular family. This varied pattern of distribution across metazoans not only challenges the conventional view of a uniform presence across animal taxa but also enhances our understanding of the intricate evolutionary dynamics that have shaped the existing diversity of g-type lysozyme presence in the animal kingdom.

### *Aplysia californica* Encodes Phylogenetically Distinct g-Type Lysozyme Genes

The sea slug *Aplysia californica* is a powerful reference species and model for understanding cellular bases of behaviors and neuroplasticity (Kandel, 1979, 2001; Moroz, 2011). As with many invertebrates, *Aplysia californica* lacks c-type or i-type lysozymes. Figure 1 highlights the presence of three g-type lysozyme genes in *Aplysia californica*, designated as LYZg1, LYZg2, and LYZg3. Notably, LYZg3 forms a unique, possible ancestral subgroup and lacks close orthologs within the Euthyneura group, which includes genera such as *Elysia, Biomphalaria*, and *Physella acuta* (Brenzinger, Schrodl, & Kano, 2021). Instead, LYZg3’s closest orthologs appear in the more distantly related Caenogastropoda clade, particularly within the genomes of *Littorina saxatilis* and *Conus magus*. The phylogenetic relationships of this gene reach even further, to distant relatives within the Patellogastropoda, such as *Lottia gigantea* and *Haliotis spp*. (Schoch et al., 2020). Due to its unique evolutionary path and distinct phylogenetic position, we have classified the more ancestral LYZg3 under a new category, termed g-type1 lysozyme genes (Figure 1).

The evolutionary analysis suggests that the acquisition of LYZg3 by *Aplysia californica* predates the more recent gene duplication events responsible for LYZg1 and LYZg2. These newer genes show closer evolutionary relationships within the Euthyneura group, with species like *Elysia chlorotica* and *Elysia crispata* each harboring two copies of similar lysozyme genes. Conversely, within the Caenogastropoda, notably in the genome of *Littorina saxatilis*, only a single copy of this gene type is present. This finding has led us to categorize LYZg1 and LYZg2 as g-type2 lysozyme genes, thus distinguishing them from the older g-type1 group (Figure 1).

Our analysis of the g-type lysozyme genes in *Aplysia californica* reveals the presence of two phylogenetically distinct gene groups: g-type1 and g-type2. A detailed sequence identity analysis showed that g-type1 genes have an average sequence identity of 68.48%, with values ranging from 44.49% to 100%, indicating higher conservation. In contrast, g-type2 genes demonstrated greater variability, with an average sequence identity of 58.28% and a range of 45.69% to 100%. These results suggest that g-type1 lysozymes may perform more conserved and essential functions. In contrast, the g-type2 lysozymes are likely subject to adaptive pressures, possibly reflecting diverse roles in immune response or digestive functions.

In addition to the sequence identity analysis, we observed that g-type1 lysozyme genes are exclusively present in the studied marine gastropods, whereas g-type2 lysozymes are found in both marine and freshwater species. This distribution suggests that g-type1 lysozymes may be optimized for the relatively stable environmental conditions of marine ecosystems, while the broader range of g-type2 lysozymes indicates that these genes may have evolved to handle the more variable and diverse environmental pressures encountered in freshwater habitats.

### Structural Conservation of g-type Lysozymes Across Metazoans

Figure 2A presents a multiple sequence alignment (MSA) of g-type lysozyme genes, including one from humans (LYG2), Anthozoan *Dendronephthya gigantea*, the more ancestral *Aplysia californica* gene (LYZg3), as well as the *Trichoplax* g-type lysozyme. Accompanying this alignment, secondary structure predictions obtained from the JPRED4 server underscore the preservation of key structural elements across these diverse taxa (Drozdetskiy, Cole, Procter, & Barton, 2015). Notably, the catalytic residues (aspartic acid and glutamic acid) are highlighted with stars on the MSA, illustrating their surprising conservation across broad phylogenetic distances. These catalytic residues are also conserved across all lysozyme types, including c-type and i-type lysozymes (Kuwano, Yoneda, Kawaguchi, & Araki, 2013; Taylor et al., 2019).

**Figure 2:**
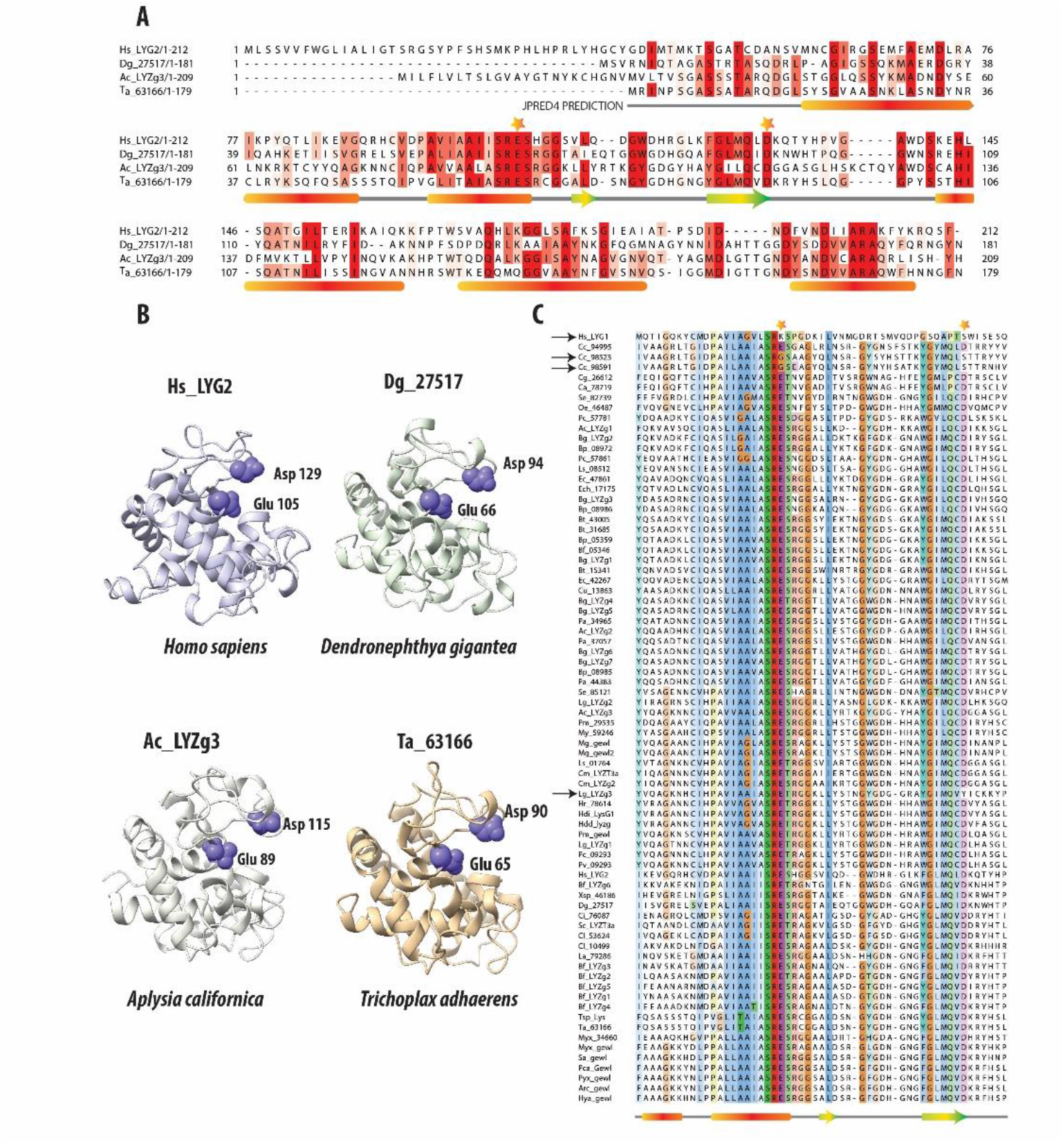
Comparative Analysis of g-Type Lysozyme Genes Across Diverse Species. *2A*: Multiple Sequence Alignment of g-type lysozyme genes from representative species, including humans, the soft coral *Dendronephthya gigantea*, one of the g-type lysozyme genes from the *Aplysia californica* genome, and a g-type lysozyme gene from the placozoan *Trichoplax adhaerens*. The alignment, enhanced by the secondary structure predictions from JPRED (Drozdetskiy et al., 2015), reveals alpha-helices marked by red bars and beta-sheets denoted by green and yellow arrows. These structural elements are essential for the enzyme’s lytic function, highlighting the evolutionary conservation despite sequence variability. Catalytic residues crucial to enzymatic activity are highlighted with stars above the alignment, emphasizing their conserved placement across species. ***2B*:** The 3D structural models of these proteins, generated by AlphaFold2 (Jumper et al., 2021), confirm the structural similarities in the regions noted in the sequence alignment. Their lysozymes also exhibit highly similar structural folds. The preservation is especially pronounced in the catalytic residues, Glutamic and Aspartic acids, which are crucial for the enzyme’s function (Malcolm et al., 1989). The consistent arrangement of these residues across representatives of different taxa underscores the structural fidelity of these proteins. It reflects evolutionary pressures to preserve key functional aspects of lysozymes crucial for antimicrobial defense. ***2C*:** Multiple sequence alignment of g-type lysozymes from a variety of species, focusing on the catalytic residues (Glutamic and Aspartic acids); these residues are marked with stars. Notably, deviations such as the non-conservation of these residues in the human Hs_LYG1 gene and two genes from *Corticium candelabrum* are indicated with black arrows. Additionally, a unique substitution in the gene Lg_LYZg3, where Tyrosine replaces Aspartic acid, is also highlighted, capturing the evolutionary dynamics of these residues.

Further, 3D structural analyses were performed using AlphaFold2 (Jumper et al., 2021). Predictions from AlphaFold2 demonstrated that the g-type lysozyme genes across a broad evolutionary spectrum fold into a structural motif observed in other lysozyme types, such as c-type or i-type (Kuwano et al., 2013). Despite lacking sequence similarity with these types, g-type lysozymes adopt similar structural conformations. Thus, the 3D structures predicted by AlphaFold2 reveal a remarkable consistency in the structural architecture of these enzymes across metazoans, again highlighting the deep evolutionary conservation of essential structural features critical for antimicrobial defense.

In order to validate the structural models of the g-type lysozyme genes across metazoans, we employed AlphaFold for structure prediction. The predicted accuracy, represented by the pLDDT (predicted Local Distance Difference Test) and pTM (predicted Template Modeling score), was calculated for each lysozyme model. As summarized in Table 1, the highest-ranked models for each gene showed consistent and high accuracy, with pLDDT values ranging from 87.4 to 97.8 and pTM values from 0.815 to 0.940, indicating reliable predictions. These results further illuminate the evolutionary preservation of g-type lysozyme architectures.

**Table 1.**
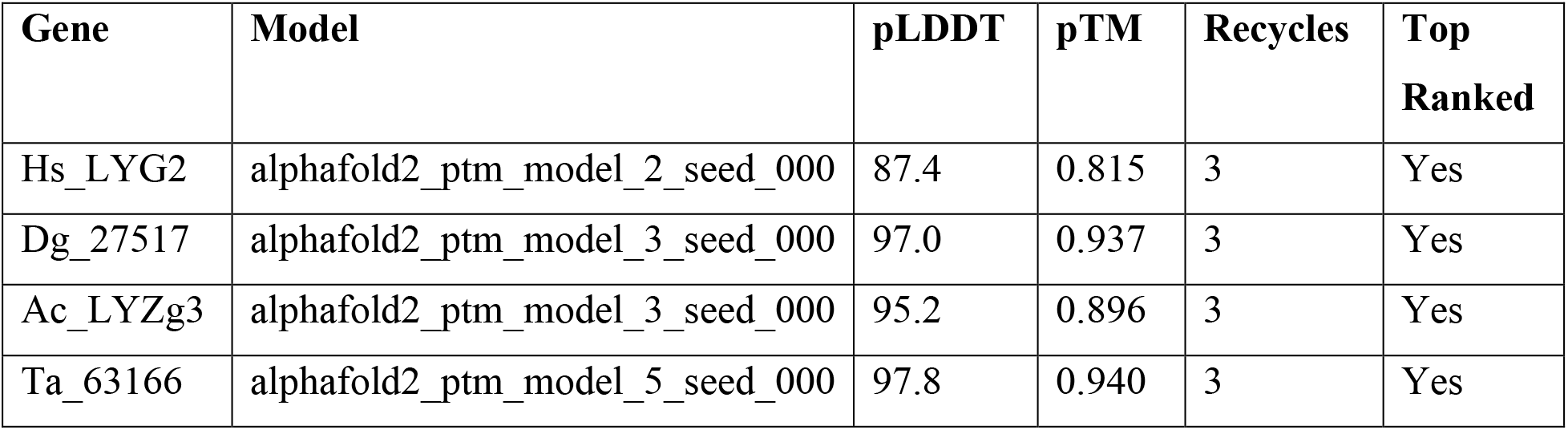
AlphaFold Model Prediction Accuracy for Selected g-type Lysozyme Genes.

In addition to the predicted accuracy, we compared the Root Mean Square Deviation (RMSD) between different model pairs for each gene, as shown in Table 2. The RMSD values across model comparisons ranged from 0.100 to 0.183, indicating close structural similarity between different AlphaFold predictions for each gene. Furthermore, the structural alignments between the AlphaFold2-predicted models and experimentally determined structures yielded the following RMSD values: 0.717 Å for Hs_LYG2, 0.566 Å for Dg_27517, and 0.503 Å for Ta_63166 (all aligned with PDB ID: 4G9S). Additionally, the alignment between the AlphaFold2-predicted model for Ac_LYZg3 and the experimentally determined structure (PDB ID: 4G9S) produced an RMSD of 0.644 Å, demonstrating a high degree of similarity between the predicted and experimentally derived conformations.

**Table 2.**
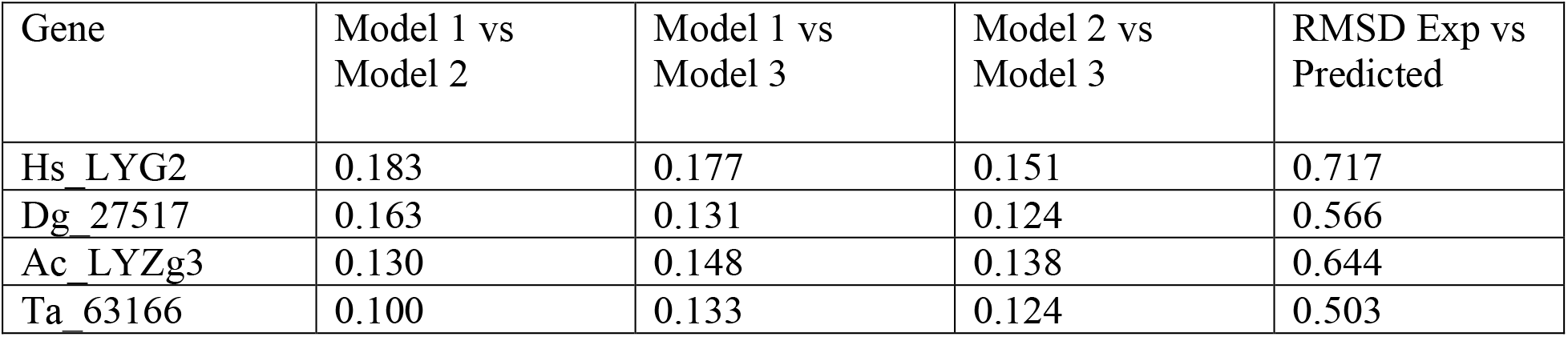
Comparison of Root Mean Square Deviation (RMSD) Between Models for g-Type Lysozyme Genes.

| Gene | Model 1 vs Model 2 | Model 1 vs Model 3 | Model 2 vs Model 3 | RMSD Exp vs Predicted |
| --- | --- | --- | --- | --- |
| Hs_LYG2 | 0.183 | 0.177 | 0.151 | 0.717 |
| Dg_27517 | 0.163 | 0.131 | 0.124 | 0.566 |
| Ac_LYZg3 | 0.130 | 0.148 | 0.138 | 0.644 |
| Ta_63166 | 0.100 | 0.133 | 0.124 | 0.503 |

### Conservation of Catalytic Residues in g-Type Lysozymes Across Metazoans

Figure 2C presents a multiple sequence alignment of g-type lysozymes from various representative species, focusing on the amino acids surrounding the critical catalytic residues, glutamic and aspartic acids. These essential residues are highlighted with stars above the alignment to underscore their importance in the enzyme’s catalytic mechanism. Previous research has demonstrated that site-directed mutagenesis of aspartic acid (D52N) and glutamic acid (E35Q) in the chicken egg white lysozyme significantly impairs enzymatic activity. The mutant enzyme D52N retains approximately 5% of the wild-type lytic activity against *Micrococcus luteus* cell walls, whereas E35Q exhibits virtually no measurable activity (0.1% ± 0.1%)(Malcolm et al., 1989). These findings affirm the pivotal role these residues play in the enzyme’s catalytic function.

Interestingly, we observed variations in the conservation of these residues across different species. In the human *Hs_LYG1* gene, both glutamic acid and aspartic acid are not conserved, as indicated by the black arrow next to the gene name. *Hs_LYG1* is highly expressed in the kidney, which plays a key role in detoxification and filtration, with lower expression levels noted in the liver and testes. A similar pattern of non-conservation is found in two of the three g-type lysozyme genes from the poriferan *Corticium candelabrum* (genes *Cc_98523* and *Cc_98591*), which also lack conservation of these catalytic residues, marked by black arrows next to the gene names. In contrast, these catalytic residues are generally conserved in most other g-type lysozymes across other taxa. An exception to this trend is the gene *Lg_LYZg3*, where the aspartic acid (D) is replaced by tyrosine (Y), again indicated by a black arrow. While the specific effects of these mutations on enzyme activity have not yet been experimentally validated, it is hypothesized that these changes may result in reduced or altered enzymatic function, as seen with the chicken lysozyme mutants (D52N and E35Q).

In summary, the broad preservation of these catalytic residues highlights their evolutionary significance in maintaining enzymatic functions, with a few notable exceptions. These exceptions likely represent lineage-specific adaptations that have altered the canonical amino acids involved in the catalytic mechanism.

### Tissue-Specific Expression Patterns of *Aplysia californica* g-Type Lysozyme Genes

Using RNA-seq data, we characterized the tissue-specific expression profiles of g-type lysozyme genes (LYZg1, LYZg2, LYZg3) with distinct expression patterns for each gene, which suggests the division of labor and specialized roles of different isoforms in physiology and development. LYZg1 and LYZg3 are associated with the digestive system and functions, whereas LYZg2 is more broadly distributed and might participate in innate immunity.

Specifically, the LYZg1 gene in *Aplysia californica* has the highest expression levels in the radula region (with a value 1061 TPM, Transcripts Per Million). The radula is equipped with chitinous radular teeth for the mechanical breakdown of food (Scheel, Gorb, Glaubrecht, & Krings, 2020). Additionally, digestive organs (esophagus and stomach) also show a significant expression (TPM of 91, Figure 3A). LYZg3 was predominantly expressed in the hepatopancreas - the major digestive gland (1610 TPM), suggesting a specialized role in food processing. This gene also showed moderate expression levels in some abdominal neurons and possible associated glia, implying immune-like functions (Figure 3E). At very low levels, LYZg1 is expressed in neural tissues (7-27 TPM), which might be associated with either non-functional ‘leakage’ of promoters in polyploid cells or some other unknown functions (Figure 3E).

**Figure 3:**
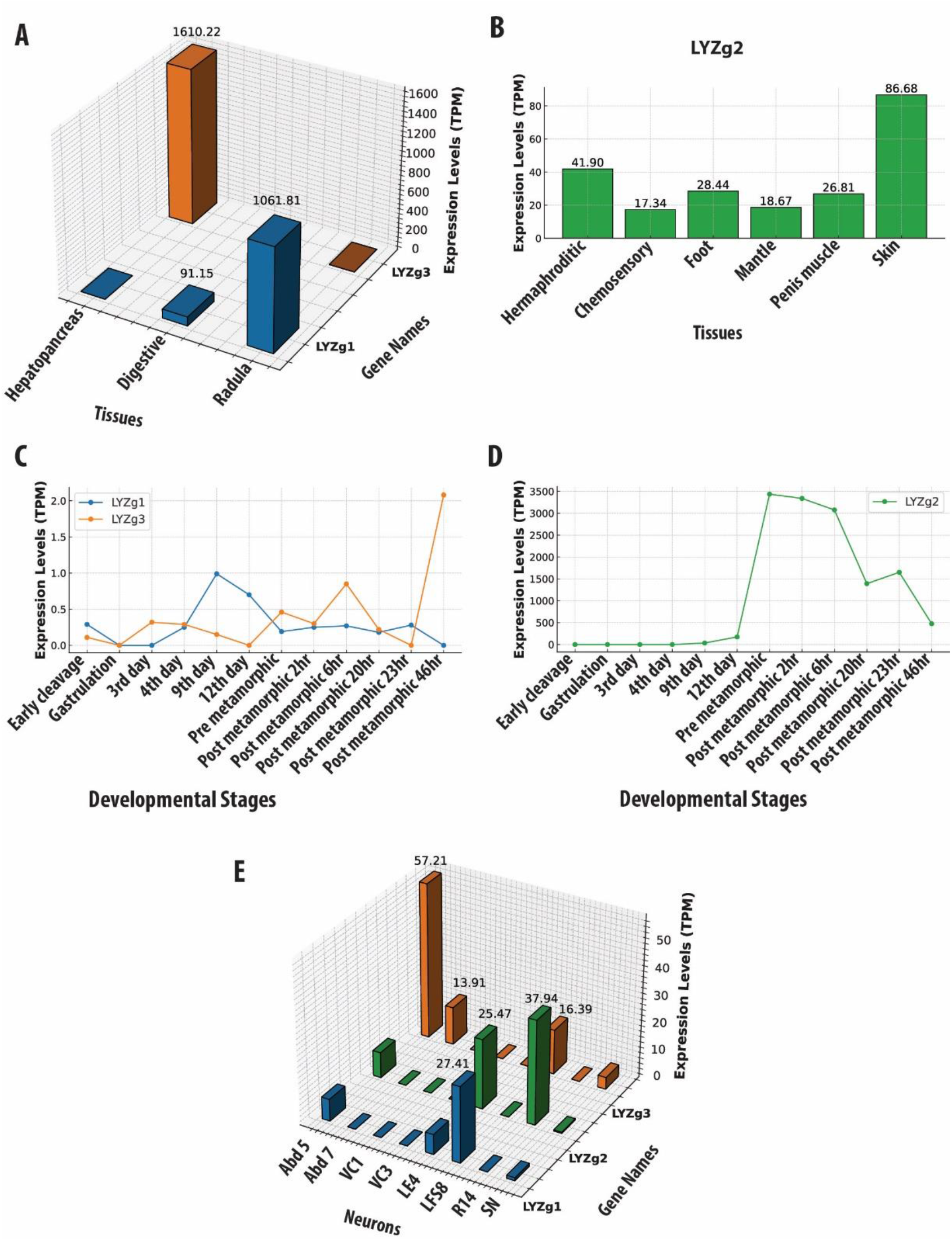
Gene Expression Profiles of *Aplysia californica* g-Type Lysozyme Genes Across Various Contexts. This figure illustrates the expression profiles of g-type lysozyme genes from *Aplysia californica* across multiple biological contexts. ***3A***: Tissue-specific expression patterns of LYZg1 and LYZg3, normalized to Transcripts Per Million (TPM). ***3B***: Tissue-specific expression patterns of LYZg2, highlighting its activity in immune-related tissues. ***3C***: Developmental stage-specific expression patterns of LYZg1 and LYZg3, showing notable peaks during early post-metamorphic stages. ***3D***: Expression levels of LYZg2 across developmental stages, indicating significant increases during post-metamorphic transitions. ***3E***: Expression profiles of LYZg1, LYZg2, and LYZg3 in individual neurons, emphasizing their roles in neural function. All expression data are normalized to TPM.

In contrast, LYZg2 showed more varied expression patterns across different tissues (e.g., the hermaphroditic glands [41TPM], chemosensory regions of mouth and rhinophores [17.31 TPM], penis muscle [26 TPM], skin [86 TPM], and foot [28 TPM] – see Figure 3B). During development, the expression of LYZg2 was noted on the 9th day (33 TPM) and significantly increased during metamorphosis, peaking in post-metamorphic stages [3339-3436 TPM [Figure 3D), suggesting its involvement in the broad spectrum of functions, during developmental and tissue differentiation, likely innate immunity. *Aplysia californica* is a simultaneous hermaphrodite with biphasic life cycle and evident metamorphosis.

## Discussion

In various organisms, lysozymes play dual roles: they protect against microbial invasions and facilitate essential digestive processes (Dobson, Prager, & Wilson, 1984; Itoh & Takahashi, 2007). This study reveals a highly mosaic distribution of g-type lysozyme genes across metazoan phyla, with the apparent absence of these enzymes in ctenophores and most sponge classes (except the homoscleromorph *Corticium candelabrum*). The potential absence of g-type lysozymes in basal groups like ctenophores and sponges may represent either their primary absence in the common metazoan ancestor or early divergence of lysozyme subtypes and gene loss in many lineages, as opposed to lateral gene transfer in bilaterians, for example, illuminated in insects (Li et al., 2022; Liu et al., 2023).

Notably, the demosponge *Suberites domuncula*, possesses a different lysozyme subtype (i-type) and employs it for immune defense and digestive processes (Wiens et al., 2005). Sponges, being bacterial feeders, utilize specialized cells, choanocytes, for a filter mechanism similar to unicellular and colonial choanoflagellates. Both sponges and choanoflagellates share a critical structural feature for feeding – a collar: an apical cilium that generates water currents to draw bacteria surrounded by microvilli, which capture bacteria for intracellular digestion by phagocytosis (Mah, Christensen-Dalsgaard, & Leys, 2014). Although we did not identify a g-type lysozyme in the sequenced choanoflagellate genomes, we discovered geneologically unrelated families of genes encoding ch-type (*Chalaropsis*-type) and other unconventional lysozyme-like proteins and their potential involvement in digestive and defensive functions.

A broader look at the distribution of g-type lysozymes across bilaterians further suggests the same two alternative hypotheses: first, these genes may be conserved across the Tree of Life, playing fundamental roles across bacteria and animals, which we think is unlikely due to the absence of these enzymes in many other eukaryotic lineages. Alternatively, the observed distribution in Metazoa could be explained by lateral gene transfer (LGT) followed by differential gene loss in some lineages, too. Additional studies exploring the phylogenetic relationships and functional similarities between bacterial and metazoan lysozymes will be necessary to pinpoint whether the pattern is a result of gene conservation or LGT events.

Placozoans, the simplest metazoans with the primary absence of neurons and muscles, have developed extracellular digestion without a mouth and gut (Romanova & Moroz, 2024). G-type lysozyme gene in *Trichoplax*, with no evidence of other lysozyme types, might likely contribute to both immunity and digestion as in soft coral species like *Dendronephthya* and *Xenia* sp., with possible functional specialization for both diet and defense. While specific physiological reports on lysozymes in anthozoans are lacking, lysozyme-like activity has been observed in the mucus of *Actinia equina* (Stabili, Schirosi, Parisi, Piraino, & Cammarata, 2015). Furthermore, our findings suggest that cnidarians, including *Nematostella vectensis*, may possess non-conventional types of lysozyme (unpublished data). The hemichordate *Saccoglossus kowalevskii* appears to encode genealogically different i-type and other non-conventional lysozyme as examples of convergent evolutionary adaptation, likely for both defensive and digestive roles.

Lysozymes broadly exhibit multifunctionality in marine bivalves, including scallops, clams, oysters, and mussels. These enzymes are utilized to degrade bacteria ensnared within the mucus of their gills, a process vital for nutrient uptake and enhancing digestive efficacy (Van Herreweghe & Michiels, 2012). Similarly, terrestrial invertebrates such as worms and flies, which feed on decomposing organic matter, depend on other families of lysozymes for both defensive measures against potential pathogens and the efficient extraction of nutrients from their microbial food sources. In flies, specifically in *Drosophila melanogaster*, lysozymes are part of the immune response system and are known to target bacterial peptidoglycans, as well as perform digestive roles in the midgut (Hultmark, 1996). Other arthropods, such as crustaceans, also utilize lysozymes for immune defense, breaking down bacterial cell walls to protect against infections (Bachali et al., 2002). In nematodes like *Caenorhabditis elegans*, lysozyme genes are expressed in the intestine, playing roles in pathogen defense and digestion (Boehnisch et al., 2011). Genealogically different i-type lysozymes also perform digestive roles akin to the function of c-type lysozymes in the foreguts of ruminants, highlighting a striking example of convergent evolution spanning diverse biological groups (Bachali et al., 2002; Olsen, Nilsen, Sletten, & Myrnes, 2003).

The observed high sequence similarity between g-type lysozymes in animals and those found in myxobacteria such as *Corallococcus, Myxococcus, Pyxidicoccus, Archangium, Stigmatella*, and *Hyalangium* presents a compelling case of both molecular convergence and horizontal gene transfer across diverse domains of life. This striking homology not only suggests that similar selective pressures may have independently shaped these enzymes in vastly different organisms but also highlights the essential functional role of lysozymes in both predation and immune defense.

In myxobacteria, these lysozyme-like proteins are likely integral to their predatory mechanisms, facilitating the breakdown of bacterial cell walls, similar to the role of g-type lysozymes in the immune systems of animals. These conserved enzymatic functions across evolutionary distances underscore the adaptive significance of lysozymes and may reflect ancient evolutionary connections (LGT) that have preserved these critical activities. This finding contributes to our broader understanding of how essential functions can be recruited or arise independently across phyla, thus offering new insights into the evolutionary dynamics of molecular function and adaptation.

The phylogenetic analysis of g-type lysozyme genes in *Aplysia californica* illuminate functional divergence and suggests distinct evolutionary pressures resulting in labor division between digestive and likely immune functions. The characteristic distribution of g-type1 and g-type2 lysozymes across marine and freshwater gastropods further supports functional divergence associated with different habitats. G-type1 lysozymes are exclusively found in marine gastropods in more stable marine environments. In contrast, g-type2 lysozymes, present in both marine and freshwater species, likely reflect phenotypic plasticity to cope with fluctuating salinity, temperature, and microbial diversity in freshwater habitats. This observation highlights shaping the evolutionary trajectories of differential recruitments of these lysozyme families, closely tied to the ecological niches. The positioning of bivalve g-type lysozyme genes between the two gastropod groups (g-type1 and g-type2) in the phylogenetic tree could reflect divergence from a common ancestral lineage shared by gastropods and bivalves or LGT events or both. This finding underscores the complexity of lysozyme evolution in molluscs.

In this context, *Aplysia californica* further demonstrates the molecular division of labor for three g-type lysozymes, showing distinct expression patterns across cells and tissues. Similarly, among scallops and abalone, g-type lysozymes are primarily located in organs vital for nutrient processing and environmental pathogen management, such as the hepatopancreas and mantle (Bathige et al., 2013). In *Aplysia californica*, LYZg1 and LYZg3 are predominantly associated with components of the digestive system, also implying that *Aplysia californica* leverages these enzymes to decompose bacterial cells, thus pursuing an evolutionary path distinct from that of mammals (Mackie, 2002). On the other hand, the pronounced expression of LYZg2 in the skin, mantle, foot, chemosensory organs, and reproductive organs likely represents *Aplysia californica* -specific adaptation, with a highly reduced shell, boosting its defenses against pathogens.

In summary, our findings highlight the multifunctional and evolutionary significance of lysozymes across Metazoa. The roles of these enzymes extend beyond immune defense, showcasing their involvement in critical digestive processes, and are shaped by lineage-specific adaptations. The observed mosaic distribution of g-type lysozymes and their varying conservation patterns point to complex evolutionary trajectories influenced by ecological and physiological pressures.

## Conclusions

1. In this study, we explored the evolutionary diversification and functional roles of g-type lysozymes across a broad range of metazoans, revealing a mosaic pattern in their distribution. Certain phyla, such as ctenophores and most sponge classes, lack these enzymes entirely, while others, like *Corticium candelabrum* and *Trichoplax*, retain or acquire g-type lysozymes that play critical roles in both immune defense and digestion. Additionally, the identification of non-conventional lysozyme-like proteins in choanoflagellates suggests that digestive and defensive functions may have evolved in parallel in these organisms despite the absence of g-type lysozymes.
2. We also uncovered significant homology between g-type lysozymes in animals and myxobacteria, indicating a possible case of molecular convergence or horizontal gene transfer. This observation highlights shared evolutionary pressures across vastly different organisms, suggesting that lysozymes may have been recruited independently throughout evolution for similar functional roles in predation and immune defense.
3. Finally, the distinct expression patterns of g-type lysozymes in various tissues of *Aplysia californica* emphasize their specialized roles. LYZg1 and LYZg3 are primarily involved in digestion, while LYZg2 plays adaptive roles in immune defense. These tissue-specific patterns reflect the evolutionary pressures that shaped lysozyme diversification and demonstrate the molecular division of labor in different metazoan lineages. There is clear divergence between g-type1 and g-type2 lysozymes in *Aplysia*. G-type1 lysozymes exhibit strong sequence conservation, likely due to their essential roles in immune defense and physiological processes. In contrast, the greater sequence variability in g-type2 lysozymes suggests adaptive evolution in response to environmental challenges, such as fluctuating microbial communities in different habitats. This functional diversification, together with the diversity of these enzymes in marine and freshwater species, highlights the role of sequence variation in shaping molluscan physiology, where g-type lysozymes serve distinct functions.

In summary, this study provides new insights into the evolutionary dynamics of g-type lysozymes and their recruitment for diverse functional roles across metazoans. Future studies, including experimental validation of lysozyme activity and further genomic analysis, will be crucial to understanding the full scope of their contributions to both immune defense and digestion.

## Methods

### 1. Sequence Retrieval and Database Preparation

We established both local and online sequence retrieval systems. Locally, we configured the NCBI standalone BLAST tool on a UNIX-based platform, accessible via the NCBI BLAST Download. We generated a dedicated BLAST database using the makeblastdb utility, incorporating genomic data across species. Additionally, we conducted searches within the Neurobase transcriptomic dataset (https://neurobase.rc.ufl.edu/) for lysozyme genes against Ctenophore transcriptomes, which we assembled using the Trinity assembler (Grabherr et al., 2011). This database enabled targeted searches for lysozyme genes using TBLASTn, with e-value thresholds set between 10^−5 and 10^−10 to ensure the identification of potential homologs with high specificity. In instances where genomic regions lacked annotated gene models, we extracted sequences surrounding predicted coding regions and subjected them to hidden Markov model analysis using FGENESH+, thereby enhancing our ability to predict gene structures accurately. We stored all the genomic information in our locally created FileMaker Pro database (http://claris.com).

We broadened our search for lysozyme genes, utilizing species-specific online platforms such as and the Ensembl genome browser. Furthermore, we employed the NCBI BLAST server to retrieve as well as cross-examine the lysozyme gene data from a wide array of sources, including metazoans, premetazoans, single-celled eukaryotes, protists, fungi, and prokaryotic origins. By maintaining the default e-value cutoffs, we achieved a comprehensive retrieval of homologous sequences, thereby considerably expanding our dataset and enriching the comparative dimension of our study. This dataset underwent rigorous manual validation and reverse BLAST searches to ensure accurate identification of lysozyme genes. Following this, we conducted multiple sequence alignments and phylogenetic reconstructions to categorize the sequences within their appropriate lysozyme groups.

### 2. Protein Domain Identification

To elucidate the structural and functional elements of the protein sequences under study, we employed a multifaceted approach for comprehensive protein domain identification. We utilized three major databases known for their robust domain detection capabilities: the Pfam database (version 34.0), the SMART database (Simple Modular Architecture Research Tool), and the NCBI’s Conserved Domain Database (CDD). Each protein sequence was processed and submitted to these search tools, which utilize hidden Markov models (HMMs) to detect known protein domains within the queried sequences. This approach allows for the identification of conserved domains that are critical to understanding the functional capabilities and evolutionary history of the proteins. The search parameters across all platforms were harmonized by setting an E-value cutoff of 0.01 to minimize the inclusion of potentially spurious matches, thereby enhancing the specificity of the domain identification process. Identified domains from Pfam, SMART, and NCBI CDD were cross-referenced to confirm domain predictions (Edgar, 2004; Letunic & Bork, 2018; Marchler-Bauer et al., 2011).

### 3. Protein Multiple-Domain Alignment

Protein sequences were aligned using the MUSCLE (Multiple Sequence Comparison by Log-Expectation) algorithm, a method recognized for its high quality and rapid alignment capabilities. We applied MUSCLE either through the European Bioinformatics Institute (EBI) online interface or directly via a command-line utility on a UNIX system. This meticulous alignment process is critical for maintaining the precision necessary for our subsequent evolutionary and functional analyses. The reliability and effectiveness of MUSCLE in generating accurate alignments make it a preferred choice in bioinformatics studies (Edgar, 2004).

### 4. Phylogenetic Analysis

We constructed phylogenetic trees using PhyML v3.0, a tool renowned for its efficiency and accuracy in tree estimation. To determine the most appropriate evolutionary model, we used ProtTest to evaluate models based on the Akaike Information Criterion (AIC). This model was applied with adjustments for rate heterogeneity and other relevant evolutionary parameters to best fit our data. Tree topology was refined employing both Nearest Neighbor Interchange (NNI) and Subtree Pruning and Regrafting (SPR) moves to optimize the tree structure. Clade support was rigorously assessed via the SH-like approximate likelihood ratio test, which provided a robust statistical framework for evaluating phylogenetic reliability and accuracy. This comprehensive approach ensures that our phylogenetic interpretations are both scientifically robust and reliable, facilitating detailed evolutionary insights (Anisimova, Gil, Dufayard, Dessimoz, & Gascuel, 2011; Guindon et al., 2010; Guindon & Gascuel, 2003).

### 5. Visualization and Post-processing of Phylogenetic Trees

Phylogenetic trees were initially visualized using iTOL (Interactive Tree Of Life), an online tool that allows for the comprehensive display and annotation of phylogenetic and other hierarchical trees. To improve visual clarity and enhance publication quality, the resulting diagrams were further refined using Adobe Illustrator. This post-processing step enabled us to fine-tune graphic details, adjust color schemes, and add text annotations, thus enhancing the overall readability and aesthetic appeal of the tree presentations. These refinements were crucial for effectively conveying complex evolutionary relationships among the studied lysozyme genes, facilitating a clearer understanding and interpretation by the scientific community. This approach ensured that our visual representations were not only scientifically accurate but also visually compelling.

### 6. Prediction of Protein Secondary and Tertiary Structures

Protein secondary structures were predicted using the JPRED4 online server, which is highly regarded for its accuracy and reliability in secondary structure prediction (Drozdetskiy et al., 2015). We uploaded the Multiple Sequence Alignment (MSA) file, saved in MSF format from the ClustalX MSA viewer, directly to JPRED4. This step facilitated the automated prediction of alpha-helices, beta-sheets, and coil regions within the protein sequences, providing a foundational understanding of the structural elements.

For the prediction of protein tertiary structures, we employed the AlphaFold2.ipynb notebook on the AlphaFold2 server hosted on Google Colab (ColabFold v1.5.5). This platform leverages MMseqs2 for sequence searching, significantly enhancing the accuracy and reliability of the protein folding predictions (Jumper et al., 2021; Mirdita et al., 2022). AlphaFold2’s deep learning-based approach allows for the generation of highly accurate 3D models of protein structures based on the amino acid sequences provided.

### 7. *Aplysia californica* cDNA Library Preparation and Sequencing

A comprehensive cDNA library for *Aplysia californica* was constructed by the Moroz laboratory as a part of the *Aplysia californica* genome/transcriptome project, using an Illumina kit, specifically designed for high-fidelity and efficient cDNA synthesis (Moroz & Kohn, 2010). The transcriptomes were sequenced using an Illumina sequencing platform. The sequencing was optimized to achieve high coverage and depth, enhancing the reliability of gene expression data.

### 8. Data Processing and Expression Analysis

The initial step involved using Trimomatic for trimming adapter sequences and removing low-quality bases, which is critical for maintaining the integrity of subsequent analyses. Following this preprocessing, the cleaned reads were aligned to the *Aplysia californica* genome using the STAR aligner, a tool renowned for its efficiency and accuracy in mapping large numbers of RNA-seq reads (Dobin et al., 2013).

After alignment, gene expression levels were quantified using EdgeR, an established bioinformatics tool for differential expression analysis (Robinson, McCarthy, & Smyth, 2010). In this phase, we focused on calculating Transcripts Per Million (TPM), a normalization method that facilitates accurate comparison of gene expression levels across different samples by accounting for both the depth of sequencing and the gene length.

### 9. Visualization of Gene Expression data

For the gene expression data from our *Aplysia californica* study, we used bar and line graphs generated in the R programming environment with the ggplot2 package. The data were log-transformed to better visualize differences in gene expression levels across tissues; however, the graphs are labeled with the actual TPM (Transcripts Per Million) values to ensure accuracy in reporting. Bar graphs were used to compare gene expression across tissues, with each bar representing the average expression in TPM, and 3D bar diagrams were employed for more complex visualizations. Line graphs were used to display temporal changes in gene expression across developmental stages or under varying experimental conditions.

## Acknowledgments

This research was supported by National Science Foundation grants (2341882) awarded to L.L.M. Additionally, this work was partly funded by the National Institute of Neurological Disorders and Stroke of the National Institutes of Health under Award Number R01NS114491 to L.L.M.

## Notes

### Competing Interest Statement

The authors have declared no competing interest.

